# On the asymptotic equivalence between the radon and the hough transforms of digital images

**DOI:** 10.1101/818799

**Authors:** Riccardo Aramini, Fabrice Delbary, Mauro C Beltrametti, Claudio Estatico, Michele Piana, Anna Maria Massone

## Abstract

Although characterized by different mathematical definitions, both the Radon and the Hough transforms ultimately take an image as input and provide, as output, functions defined on a preassigned parameter space, i.e., the so-called either Radon or Hough *sinograms*. The parameters in these two spaces describe a family of curves, which represent either the integration domains considered in the Radon transform, or the kind of curves to be detected by the Hough transform.

It is heuristically known that the Hough sinogram converges to the corresponding Radon sinogram when the discretization step in the parameter space tends to zero. By considering generalized functions in multi-dimensional setting, in this paper we give an analytical proof of this heuristic rationale when the input grayscale digital image is described as a set of grayscale points, that is, as a sum of weighted Dirac delta functions. On these grounds, we also show that this asymptotic equivalence may have a valuable impact on the image reconstruction problem of inverting the Radon sinogram recorded by a medical imaging scanner.

## 1. Introduction

The Radon transform (RT) [1] is an important tool in harmonic analysis and inverse problems theory with significant impacts on both group theory and applied mathematics. The classical definition of this transform considers integrals over hyperplanes with specific orientation and distance from a reference hyperplane. In this case, many functional properties of the RT have been investigated, including the characterization of its kernel and range, the ill-posedness of the inverse problem, as well as several inversion formulas and algorithms. The RT has also been extended to integration on manifolds [2], although here important functional and computational problems are still open. In biomedical imaging, RT is the well-established mathematical model for data formation in X-ray Computerized Tomography (CT) and in Positron Emission Tomography (PET) [3, 4] and all software tools for image visualization implemented in current industrial CT and PET scanners must realize, at same stage, the numerical inversion of RT.

While RT plays a crucial role in image reconstruction, the Hough transform (HT) [5, 6, 7] provides an important computational technique in pattern recognition. Indeed, HT is widely used in image-processing to detect specific algebraic plane curves, which are zero-loci of polynomials whose coefficients depend polynomially on a (small) set of parameters. The basic idea of this recognition procedure comes from the point-line duality in projective plane geometry. Indeed, any single point in the image space corresponds to a locus (which is its HT) in the parameter space, so that each parameter in the locus identifies a curve passing through the point. In turn, a curve in the image space corresponds to a single point in the parameter space, given by the intersection of all HTs of the points belonging to the curve. A histogram (the Hough counter) can therefore be constructed, representing an accumulator function defined on the discretized parameter space: for each cell in the discretized parameter space, the value of the accumulator corresponds to the number of HTs passing through that cell. This way, the position of the maximum in the Hough counter identifies the set of parameters characterizing the curve to be detected in the image space. Improved algorithms based on HT voting schemes and enabling very rapid and efficient recognition of specific geometric features in large images or data sets, have been introduced in [8, 9]. From a more theoretical point of view, a recent research [10] provided a rigorous mathematical foundation, based on algebraic-geometry arguments, for the case of algebraic plane curves of whatever degree, together with a key lemma stating equivalent conditions under which the existence and uniqueness of the intersection point of the HTs is guaranteed. This framework was then applied in [11, 12, 13, 14] to provide an atlas of algebraic curves used to recognize profiles in real astronomical and biomedical images.

This paper formally investigates a known relationship between RT and HT which has been so far discussed just from a heuristic perspective. Specifically, our aim is to prove analytically that, given a digital image represented as a sum of weighted Dirac delta functions, the corresponding Hough counter tends to become the RT of the image itself as the discretization of the parameter space becomes finer and finer. The basic idea dates back to 1981, when an IEEE letter [15] provided examples in which HT can be considered as a particular case of RT. Although influential and constructive, this letter is heuristic and does not consider any formal definition of HT. About ten years later, in [16], the limitations of [15] were noticed and the similarity between the two transforms was furtherly investigated by relying on such a formal definition. Later on, in 2004, an inspirational report [17] outlined a sort of research program to properly understand the link between the two computational tools. However, none of these studies provided a consistent and rigorous proof of the convergence of the Hough counter to the RT sinogram.

The present paper aims to fill this gap setting up a general formalism for a unified description of the two transforms based on well-established mathematical tools, providing a general theorem proving the asymptotic superposition of the Radon and Hough sinograms, and showing a first hint about how this result can be utilized in image reconstruction.

The structure of the paper is as follows. Section 2 provides a summary of the formalism necessary to prove the convergence theorem, which is illustrated in Section 3. Section 4 shows a first application of the theorem to image processing. Our conclusions are offered in Section 5.

## 2. The Radon Transform and the Hough Transform

This section provides an overview of basic definitions and results concerning RT and HT. The level of generality we adopted is coherent with the formal framework necessary to prove the convergence theorem discussed in Section 3, which is the core result of this paper.

### 2.1 The Radon Transform

The RT of a piecewise continuous function with compact support in ℝ^*n*^ can be defined over a (*n* − 1)-dimension manifold in an open subset of ℝ^*n*^, parameterized by a finite number *t* of parameters varying on a open subset of ℝ^*t*^, according to the following.

#### Definition 1.

*For n* ∈ ℕ \{0, 1} *and t* ∈ ℕ \{0}, *let W and E be non-empty open subsets of* ℝ^*n*^ *and* ℝ^*t*^ *respectively, and let E*′ := {*λ*′ ∈ ℝ^*t-*1^ : ∃*λ*_*t*_ ∈ ℝ : *λ* = (*λ*′, *λ*_*t*_) ∈ *E*} *(with E*′ = Ø *if and only if t* = 1*). Moreover, let f* : *W* × *E* → ℝ *be a function expressible in the λ*_*t*_*-solvable form, i*.*e*., *as f* (*x*; *λ*) := *λ*_*t*_ - *F* (*x*; *λ*′), *being F* : *W × E*′ → ℝ *such that F* ∈ *C*^1^ (*W × E*′), *and assume that*

i. *𝒮*(*λ*) := {*x* ∈ *W* : *f* (*x*; *λ*) = 0} ≠ Ø ∀*λ* ∈ *E;*
ii. grad*x f*(*x*; *λ*) ≠ 0 ∀*λ* ∈ *E*, ∀*x* ∈ *𝒮*(*λ*).

*Finally, let m be a piecewise continuous function with compact support defined on W, and denoted by* 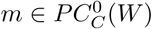. *Then, the* generalized Radon transform (GRT) *of m is the function* (*R*_*f*_ *m*) : *E* → ℂ *defined by*

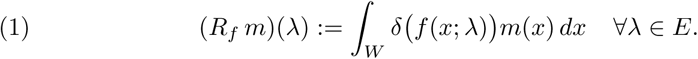

Here *δ*(*f* (·; *λ*)) denotes the *δ*-Dirac distribution of the function *f*(·; *λ*) : *W* → ℝ, associated to the submanifold 𝒮(*λ*) ⊂ *W*, ∀*λ* ∈ *E* [18]. The *λ*_*t*_-solvability condition for *f* in this definition is not necessary for the following results to hold, but we assume it since it will notably simplify the formalism.

It is also possible to define the GRT of a *δ*-Dirac distribution by first recalling that any locally integrable function 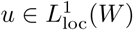 uniquely defines a corresponding distribution 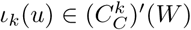, that is a continuous linear functional 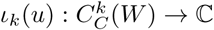, by setting

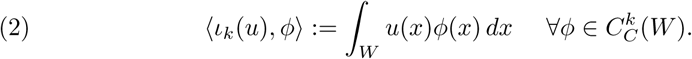

Here, for any *k* ∈ ℕ or 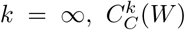 denotes the space of compactly supported *k*-times continuously differentiable functions or the space of compactly supported infinitely differentiable functions, respectively, defined on *W* and, accordingly, 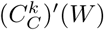 denotes the corresponding linear space of distributions. In this respect, 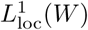 can be identified as a subspace of 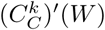, and then consider *ι*_*k*_ as an inclusion map, i.e.,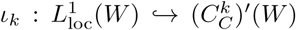. Moreover, we recall that any distribution 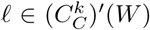 admits (distributional) partial derivatives 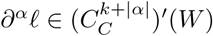 of any order |*α*| ≥ 0, according to the following

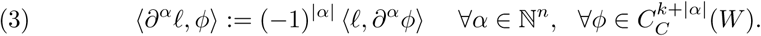

Then, the following theorem holds.

#### Theorem 1

*(see* [19] *and lemma 2*.*1 in* [20]*) Let W*, 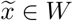 *and K* ⊂ *W be a non-empty open subset of* ℝ^*n*^, *a given point of W and a compact subset of W containing a neighborhood* 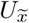 *of* 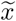, *respectively. Moreover, let* 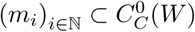 *be sequences of functions such that* supp *m*_*i*_ ⊂ *K* ∀*i* ∈ ℕ *and* 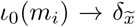 *in* 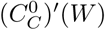 *as i* → *∞ i*.*e*.,

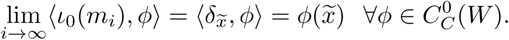

*Then, it holds that* 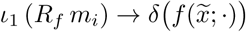 *in* 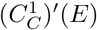 *as i* → *∞, i*.*e*.,

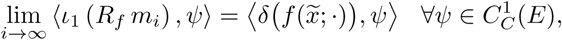

*where* 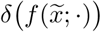 *is the Dirac delta of the function* 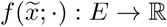.

From Theorem 1 it follows that the appropriate definition of the GRT of a single Dirac delta 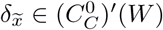 is

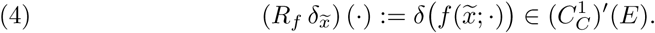

In addition, the specific form of (4) for the (classical, i.e., not the generalized) Radon transform (RT) on hyperplanes *𝒫*(*ω, γ*) := {*x* ∈ ℝ^*n*^ : *γ - ω* · *x* = 0}, ∀(*ω, γ*) ∈ (ℝ^*n*^ *\*{0}) × ℝ, becomes

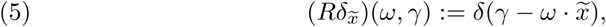

where 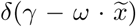 is the Dirac delta 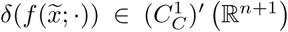 of the function mapping (*ω, γ*) into 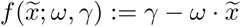.

In the following we will model a digital image *m* as the sum of a finite number *ν* of *δ*-Dirac functions, each one centered in the pixel center *x*(*P*_*j*_) and with amplitude *µ*_*j*_ ≥ 0 equal to the pixel content, ∀*j* = 1, … *ν*, i.e.

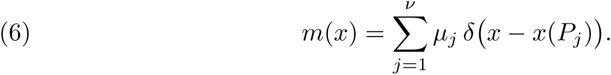

Therefore, by linearity the classical RT of *m* is

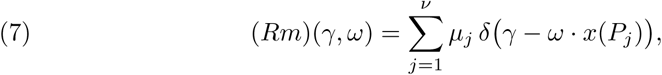

and its GRT is

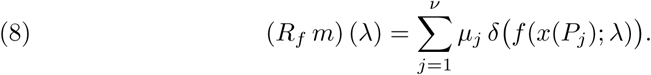

The visual representations of the intensity values of RT in (7) and of GRT in (8), with respect to the coordinates *λ* ∈ *E*, are called sinogram and generalized sinogram, respectively.

### 2.2. The Hough Transform

The Hough transform (HT) is a pattern recognition technique for the automated detection of shapes (such as lines, curves or surfaces) in images [5, 7, 10, 16, 21, 22]. In our framework an appropriate definition of HT is as follows.

#### Definition 2

*Let f* : *W* × *E* ⊂ ℝ^*n*^ × ℝ^*t*^ → ℝ *be a function such that, for each λ* ∈ *E, the map f*_*λ*_ := *f* (·; *λ*) : *W* → ℝ *defined by x* ↦ *f* (*x*; *λ*) *satisfies the following three conditions*

a. *𝒮*(*λ*) := {*x* ∈ *W* : *f*_*λ*_(*x*) = 0} ≠ Ø;
b. *f*_*λ*_ ∈ *C*^1^(*W*);
c. (grad*x f*_*λ*_)(*x*) ≠ 0, ∀*x* ∈ *𝒮*(*λ*).

*Let P* ∈ *W be a point in the image space having coordinates x* = (*x*_1_(*P*), …, *x*_*n*_(*P*)), *and let f*_*x*_ : *E* → ℝ *be the map defined by λ ↦ f*(*x*; *λ*). *Then we say that the zero locus of f*_*x*_, *defined as ℋ*(*x*) := {*λ* ∈ *E* : *f*_*x*_(*λ*) = 0} ⊂ *E, is the* Hough transform (HT) of the point *P* with respect to the function *f. If no confusion will arise, we simply say that ℋ*(*x*) *is the* HT of *x*.

By considering *𝒮*(*λ*) = {*x* ∈ *W* : *f*_*λ*_(*x*) = 0} *⊂ W* and *ℋ*(*x*) = {*λ* ∈ *E* : *f*_*x*_(*λ*) = 0} ⊂ *E*, for each (*x, λ*) ∈ *W* × *E*, the following *duality condition* [10] holds true:

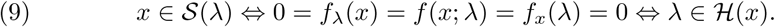

This allows us to conclude that *the* HT ℋ(*x*) *of a point x* ∈ *W contains a point λ* ∈ *E if and only if 𝒮*(*λ*) *passes through x*.

In the case of a digital image, a histogram, called weighted Hough counter (or weighted Hough accumulator), is constructed according to the following three steps.

#### I. Discretization of the parameter space

Identify a suitable (and bounded) investigation domain *𝒯* ⊂ *E* in the parameter space ℝ^*t*^. Next, choose an *initialization point* 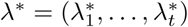 in *𝒯* and, for each *k* = 1, …, *t*, a *sampling distance d*_*k*_ with respect to the component *λ*_*k*_. Then, set

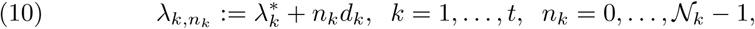

where 𝒩_*k*_ is the number of considered samples for such component, and *n*_*k*_ the index labelling the sample. Moreover, denote by

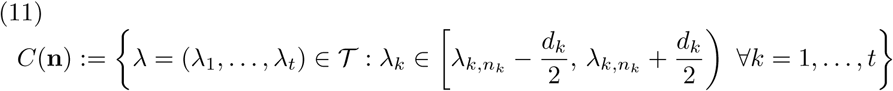

the rectangular cell with centre in the *sampling point* 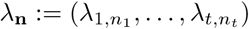 of the discretized region 𝒯, where **n** ∈ ℕ^*t*^ denotes the multi-index (*n*_1_, …, *n*_*t*_) labelling the cells. Finally, denote by *C*(*λ*) the cell containing the point *λ* ∈ 𝒯. We point out that the discretization defined by (10) only depends on the choice of the initialization point *λ*\* ∈ 𝒯 and the *discretization step* vector 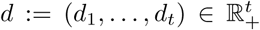. In the following, such a discretization will be denoted by {*λ*\*, d}.

#### II. *Definition of the* HT *kernel*

Similarly to Definition 1, we now focus again on a *λ*_*t*_-solvable function *f* : *W* × *E* → ℝ, that is, a function *f* satisfying (a)-(b)-(c) of Definition 2 and the following two conditions:

(d) *f* is *λ*_*t*_-*solvable*, i.e., *f* (*x*; *λ*) is of the form *f* (*x*; *λ*_1_,…, *λ*_*t*_) = *λ*_*t*_*-F* (*x*; *λ*_1_,…, *λ*_*t-*1_);
(e) for each *x* ∈ *W*, the function *f*_*x*_ : *E* → ℝ introduced in Definition 2 is continuously differentiable, i.e., in view of (d), *F*_*x*_ ∈ *C*^1^(*E*′), where, for each *x* ∈ *W*, the map *F*_*x*_ : *E*′ → ℝ is obviously defined as *λ*′ *↦ F* (*x*; *λ*′).

To introduce the Hough kernel, we first consider the function *c*_*k*_ mapping the *k*-th parameter *λ*_*k*_, for each *k* = 1,…, *t*, to its closest discretized value, which, according to (10) and (11), is defined as 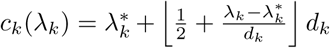, where ⌊·⌋ denotes the floor function. Moreover, by setting *𝒯*^′^ := *λ*′ ∈ ℝ^*t-*1^ : ∃*λ*_*t*_ ∈ ℝ : (*λ*′ *λ*_*t*_) ∈ *𝒯*, we also define the map *c*′ : *𝒯*′ → ℝ^*t-*1^ as *λ*′ = (*λ*_1_,…, *λ*_*t-*1_) ↦ *c*′(*λ*′) = (*c*_1_(*λ*_1_),…, *c*_*t-*1_(*λ*_*t-*1_)). Then the Hough kernel is the map *p*(·, · ; *λ*\*, *d*) : *W × T* → {0, 1}, depending on both the function *f* and the discretization {*λ*\*, *d*} of (10)–(11), defined as

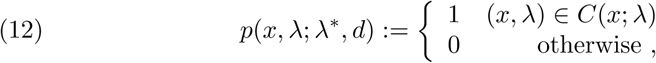

where *C*(*x*; *λ*) := (*x, λ*) *W*× *𝒯*: −*d*_*t*_*/*2 ≤ s*λ*_*t*_ *F*(*x*; *c*′(*λ*′)) < *d*_*t*_*/*2}.

#### III. Construction of the weighted Hough accumulator

For any given set of points of interest in the image space, say *P*_*j*_, *j* = 1,…, *ν*, denote by *µ*_*j*_ the grey level [23] associated with *P*_*j*_. Let us introduce the *weighted Hough counter (with respect to the function f)*, also called *weighted Hough accumulator*, as the map *H*(·; *λ*\*, *d*) :*𝒯* → ℕ defined by

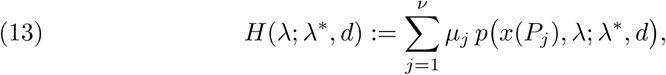

where *x*(*P*_*j*_) ∈ ℝ^*n*^ denotes the *n* coordinates of the point *P*_*j*_. Accordingly, we can now define the so-called rescaled Hough counter, which is the basis of the link between the HT and the GRT investigated in the following section.

##### Definition 3

*Let f* : *W* × *E* → ℝ *be a function satisfying conditions* (a)–(e) *of the step* II *above, and let* 𝒯 ⊂ *E be the investigation domain introduced in item* I. *Then, the function mapping λ* ∈ 𝒯 *to H*(*λ*; *λ*\*, *d*)*/d*_*t*_ *(i*.*e*., *the ratio between the weighted Hough counter defined in* (13) *with HT kernel* (12) *and the sampling distance d*_*t*_ *with respect to the component λ*_*t*_*) will be called* rescaled (weighted) Hough counter. *A visual representation of the intensity values of H*(*λ*; *λ*\*, *d*)*/d*_*t*_ *in the coordinate system* (*λ*_1_, …, *λ*_*t*_) *will be called* rescaled Hough sinogram.

## 3. The convergence theorem

The main result of the present paper is the following

### Theorem 2

*Let f* : *W* × *E* → ℝ *be a function satisfying the properties* (a)–(e) *already assumed in the previous two sections, i*.*e*., (a) 𝒮(*λ*) := {*x* ∈ *W* : *f*_*λ*_(*x*) = 0} ≠ Ø ∀*λ* ∈ *E;* (b) *f*_*λ*_ ∈ *C*^1^(*W*) ∀*λ* ∈ *E;* (c) (grad_*x*_ *f*_*λ*_)(*x*) ≠ 0 ∀*x* ∈ 𝒮(*λ*), ∀*λ* ∈ *E;* (d) *f* (*x*; *λ*_1_, …, *λ*_*t*_) = *λ*_*t*_ − *F* (*x*; *λ*_1_, …, *λ*_*t*−1_); (e) *F*_*x*_ ∈ *C*^1^(*E′*) ∀*x* ∈ *W. Moreover, let* {*λ*\*, *d*} = {*λ*\*, (*d*_1_, …, *d*_*t*_)} *be a discretization of the parameter space as explained in item* I *of the previous section, and define D* := max{*d*_1_, …, *d*_*t*_}. *Finally, let m be a discrete image (6)*, (*R*_*f*_ *m*)(*λ*) *its generalized Radon transform GRT (8), and H*(*λ*; *λ*\*, *d*)*/d*_*t*_ *the corresponding rescaled Hough counter of Definition 3 with HT kernel (12), defined on a bounded and open investigation domain* 𝒯 ⊂ *E*.

*Then the rescaled Hough counter tends to the GRT as the discretization step tends to zero, that is*

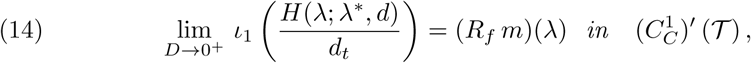

*where* 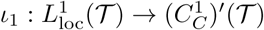 *is the inclusion map* (2).

*Proof*. We first consider the simplest digital image *m*, i.e., a single Dirac delta function located at a fixed point *P* ∈ *W*. Then, from linearity properties, the generalization to a digital image *m* of kind (6) will straightforwardly conclude the proof.

In the case of a single delta function, the statement of the theorem becomes:

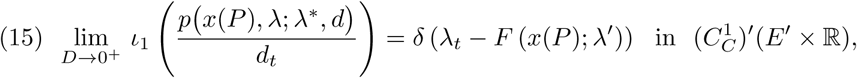

where *p* is the HT kernel (12) used in the rescaled Hough counter of Definition 3. According to (12) and (2), the thesis (15) can be recast as

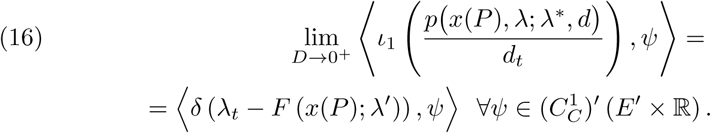

Note that 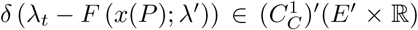 is well-defined because, for *λ* = (*λ ′, λ*_*t*_) ∈ *E′* × ℝ, we have [18]

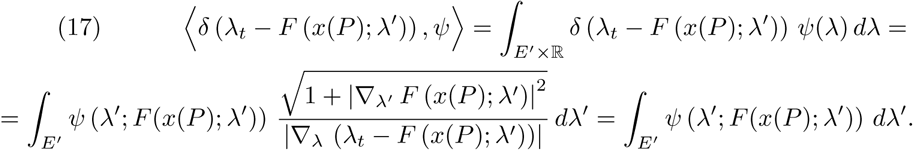

In order to prove (16), by considering (2) we have

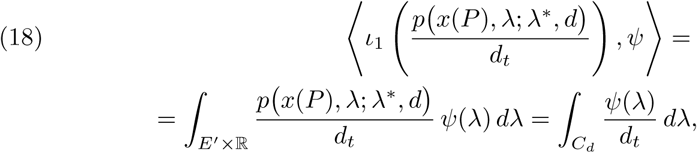

where

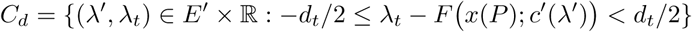

is the set of discretized cells uniquely determined by the submanifold *λ*_*t*_−*F x*(*P*); *λ′*) = 0. The last integral of (18) has a key role in our proof. Indeed, its integration domain *C*_*d*_ is a multidimensional strip, in general non continuous, contained in *E* ⊂ ℝ^*t*^, whose width with respect to the coordinate *λ*_*t*_ is *d*_*t*_, which is also equal to the denominator of the integrand. This means that, as *D* goes to zero, hence *d*_*t*_ too, the value of the integral tends, locally, to the value of *ψ* on the submanifold of the parameter space *E* such that *λ*_*t*_ *F x*(*P*); *λ′*) = 0, by virtue of Lagrange mean value theorem. The following part of the proof actually gives a mathematical evidence to this rationale. On these grounds, first we set

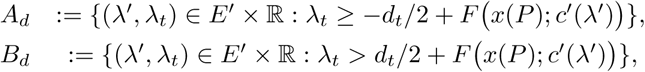

so that we can write

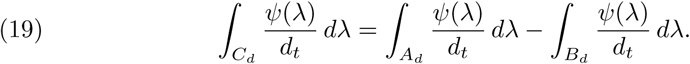

Both these two integration domains depend on the discretization step *d*. Since our aim is to relate these domains with the zero loci of *f* (which does not depend on the discretization step *d*), we formally consider the shift of the *t*-th coordinate as follows

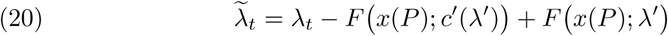

so that we can rewrite the two domains with respect to this new coordinate as

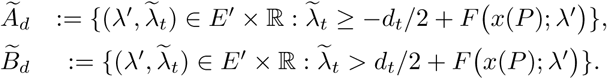

Hence, since the determinant of the Jacobian of any pure shift coordinate transformation is 1, we can write

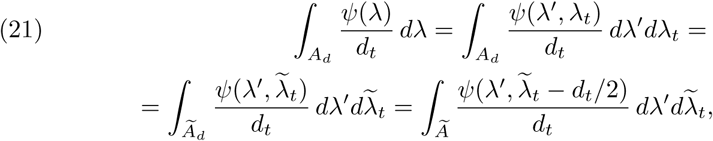

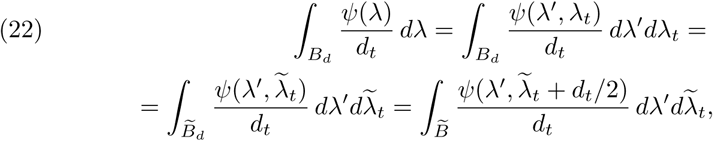

where the two sets

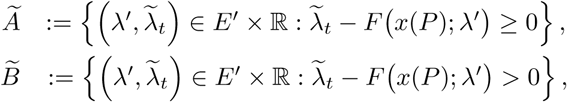

coincide up to a zero-measure subset of *E*′ × ℝ, so that we could use as well the domain *Ã* instead of 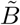 in the last integral. Accordingly, from (18) and (19),

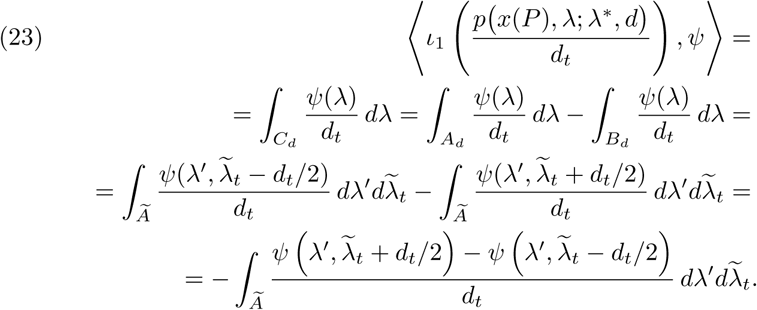

Thus, by Lagrange theorem, there exists 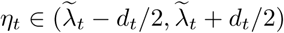 such that

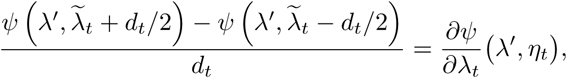

so that, by the continuity of *∂ψ*/*∂*λ_*t*_ and of *F* and the fact that 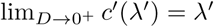 we find

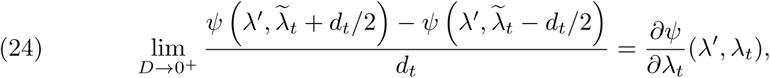

recalling that, since 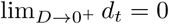, then 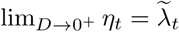 and 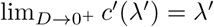, so that 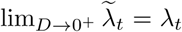. This way, by considering an upper bound 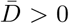 for the maximum sampling distance *D*, since 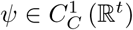, we have

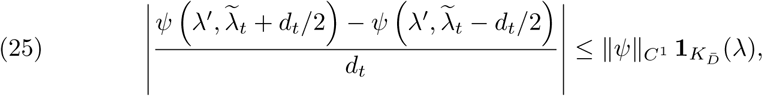

where 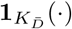 denotes the characteristic function of the compact subset of ℝ^*t*^ defined as 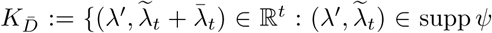 supp ψ and 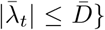. Because 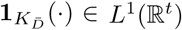, by (24)–(25) we can apply Lebesgue dominated convergence theorem in (23), thus obtaining

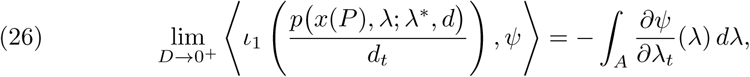

where

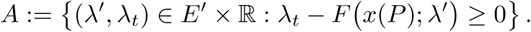

The right hand side of (26) can be rewritten by means of the strict relationships between derivative of a Heaviside step function and the Dirac delta functional. Generally speaking, given a real function *g* ∈ *C*^1^(*E*), let Θ(*g*) : *E* → {0, 1} be the characteristic function of the region {*λ* ∈ *E* : *g*(*λ*) ≥ 0}, i.e.,

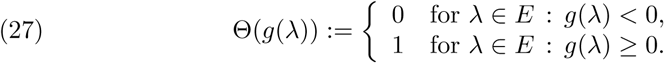

Obviously, Θ(*g*) = Θ ○ *g*, where Θ is the Heaviside function, i.e., Θ(*t*) = 0 for *t* < 0 and Θ(*t*) = 1 for *t* ≥ 0. Note that 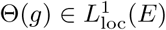: then, recalling the inclusion map 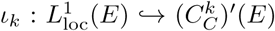 defined by (2), Θ(*f*) can also be regarded as an element of 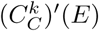, i.e., as *ι*_*k*_(Θ(*f*)), which admits partial derivatives, in a distributional sense, with respect to *λ*_*i*_, for all *i* = 1, …, *t* [18]. In particular, for any *k* ∈ ℕ or *k* = ∞, by chain rule it can be proved that its partial derivatives are as follows

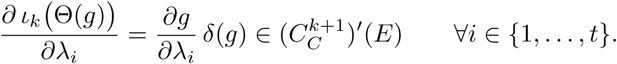

Hence, for *g*(*λ*) = *λ*_*t*_ - *F* (*x*(*P*); *λ*′), since the characteristic function of the domain *A* can be written as ℝ^*t*^ ∋ λ = (λ′, λ_*t*_) ↦ Θ (λ_*t*_ - *F* (*x*(*P*); λ′)), by considering (3) with *ℓ* = *ι*_0_ (Θ (*λ*_*t*_ - *F* (*x*(*P*); λ′))), *W* = *E, n* = *t, ϕ* = *ψ* and *α* = (0, …, 0, 1), we finally have

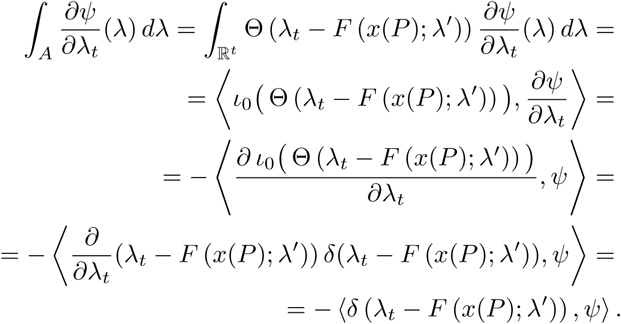

Now, a straightforward comparison between the two last equalities proves (16), equivalent to (15).

Consider now a discrete image (6) as a weighted sum (or linear combination) of Dirac delta functions. From above we have that

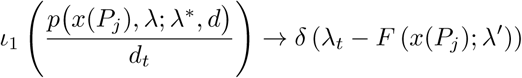

in 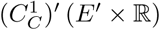 as *D* → 0^+^, for each *j* = 1, …, *ν*. The limit

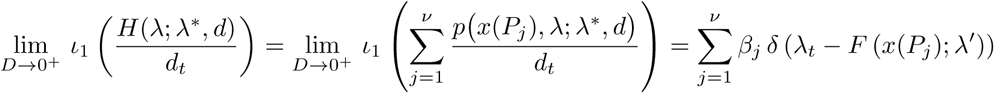

immediately follows from the linearity of the inclusion map 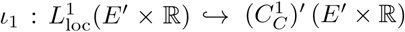. Finally, it suffices to recall that the right-hand side of the last equalities is just the GRT of *m* as shown in (8). Accordingly, this is equivalent to relation (14), thus proving the theorem. □

## 4. Numerical applications

The theoretical result contained in Theorem 2 may have significant consequences on imaging applications. Just as an example, this section shows that dealing with a noisy Radon sinogram as a (rescaled) Hough sinogram allows a more accurate image reconstruction, particularly when the noise level affecting the data is high. We consider the well-known Shepp–Logan phantom [24], shown in panel (a) of Figure 1, as the piecewise constant image (contained in a square of sides ranging from 1 to 1 and formed by pixels with values ranging from 0 to 1) to be recovered from a very noisy Radon sinogram. To this end, we first compute the exact RT with respect to the family of straight lines of equation *γ-x*_1_ cos *ϑ-x*_2_ sin *ϑ* = 0, which is of the form *f* (*x*; *λ*) = 0, under the identifications *x* = (*x*_1_, *x*_2_), *λ* = (*λ*_1_, *λ*_2_) = (*ϑ, γ*) and *f* (*x*; *λ*) = *λ*_2_ − *x*_1_ cos *λ*_1_ − *x*_2_ sin *λ*_1_. Such computation can be easily performed as explained in [25], by considering *I* discretized values *ϑ*_*i*_ (with *i* = 1 …, *I*) of *ϑ* ∈ [0, *π*) and *J* discretized values *γ*_*j*_ (with *j* = 1, …, *J*) of 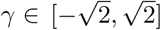. The corresponding noise-free sinogram, obtained for *I* = 629 and *J* = 287, is represented in the upper part of panel (b) in Figure 1.

**Figure 1.**
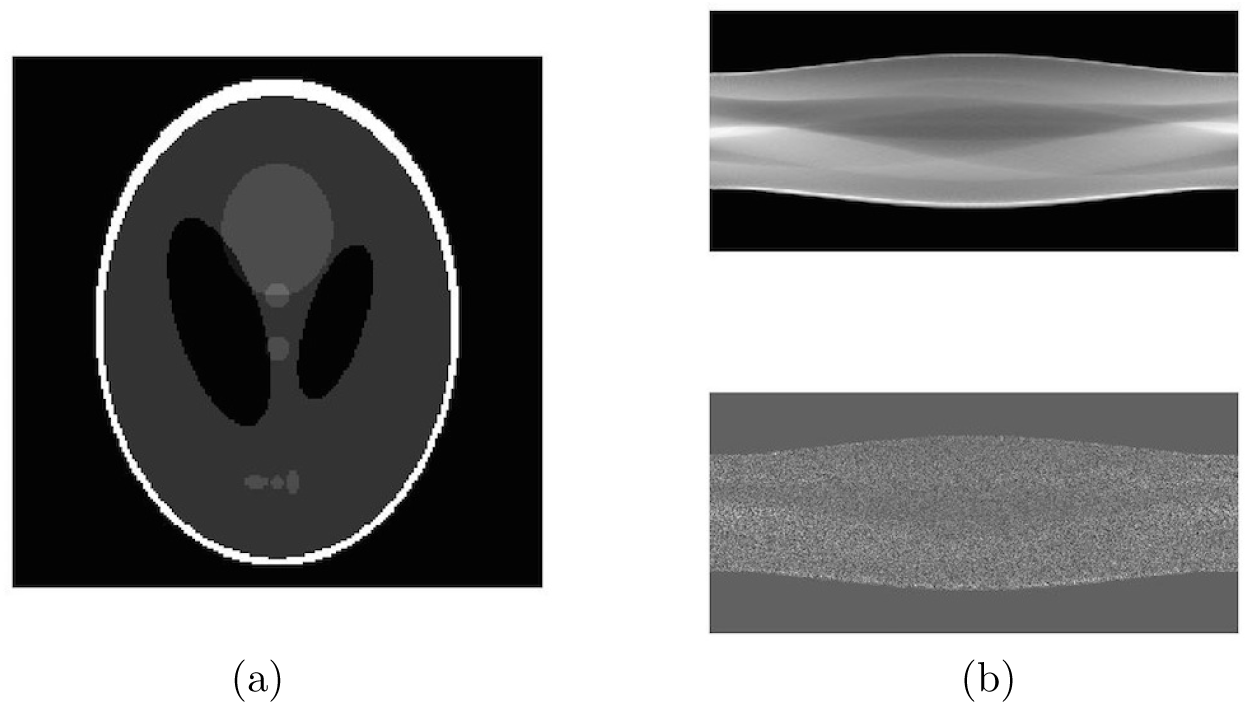
(a) The Shepp–Logan phantom. (b) Upper plot: the noise-free Radon sinogram of the Shepp–Logan phantom, computed for 629 values of *ϑ* ∈ [0, *π*) (horizontal axis) and 287 values of 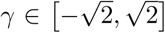 (vertical axis). Lower plot: the noisy Radon sinogram of the Shepp–Logan phantom, obtained from the upper one by corrupting it with additive 100% level Gaussian noise.

The noise-free sinogram is then corrupted by additive Gaussian noise by using the formula *S*_*n*_(*ϑ*_*i*_, *γ*_*j*_) = *S*_*t*_(*ϑ*_*i*_, *γ*_*j*_) + *𝓁 ε S*_*t*_(*ϑ*_*i*_, *γ*_*j*_), where *S*_*t*_(*ϑ*_*i*_, *γ*_*j*_) is the true value of the sinogram at the point (*ϑ*_*i*_, *γ*_*j*_), *S*_*n*_(*ϑ*_*i*_, *γ*_*j*_) is the noisy value of the sinogram at the point (*ϑ*_*i*_, *γ*_*j*_), *ε* is a realization of a normal Gaussian random variable, and *𝓁* is the relative noise level (here *𝓁* = 100% is always assumed). The resulting noisy Radon sinogram is shown in the lower part of panel (b) in Figure 1.

Usually, the inversion of RT is numerically performed by using the filtered back-projection (FBP) algorithm, where the presence of a ramp filter (Ram–Lak filter) in the frequency domain attenuates the blurring effect of a naive unfiltered back-projection and where, at the same time, a second filtering function multiplying the ramp filter allows the attenuation of high frequency noise that can be present in the Radon sinogram. Common choices for this filtering function are [26, 4] the Shepp–Logan filter; the Cosine filter; the Hamming window; the Hann window. We have then applied both the unfiltered and the filtered back-projection to recover the Shepp–Logan image from its noisy Radon sinogram, using all cited filters in the case of the FBP algorithm. The corresponding results are shown in Figure 2. It is clear that, independently of the particular filter adopted, the FBP algorithm fails to recover the internal structure of the phantom, while the unfiltered back-projection can visualize its main features.

**Figure 2.**
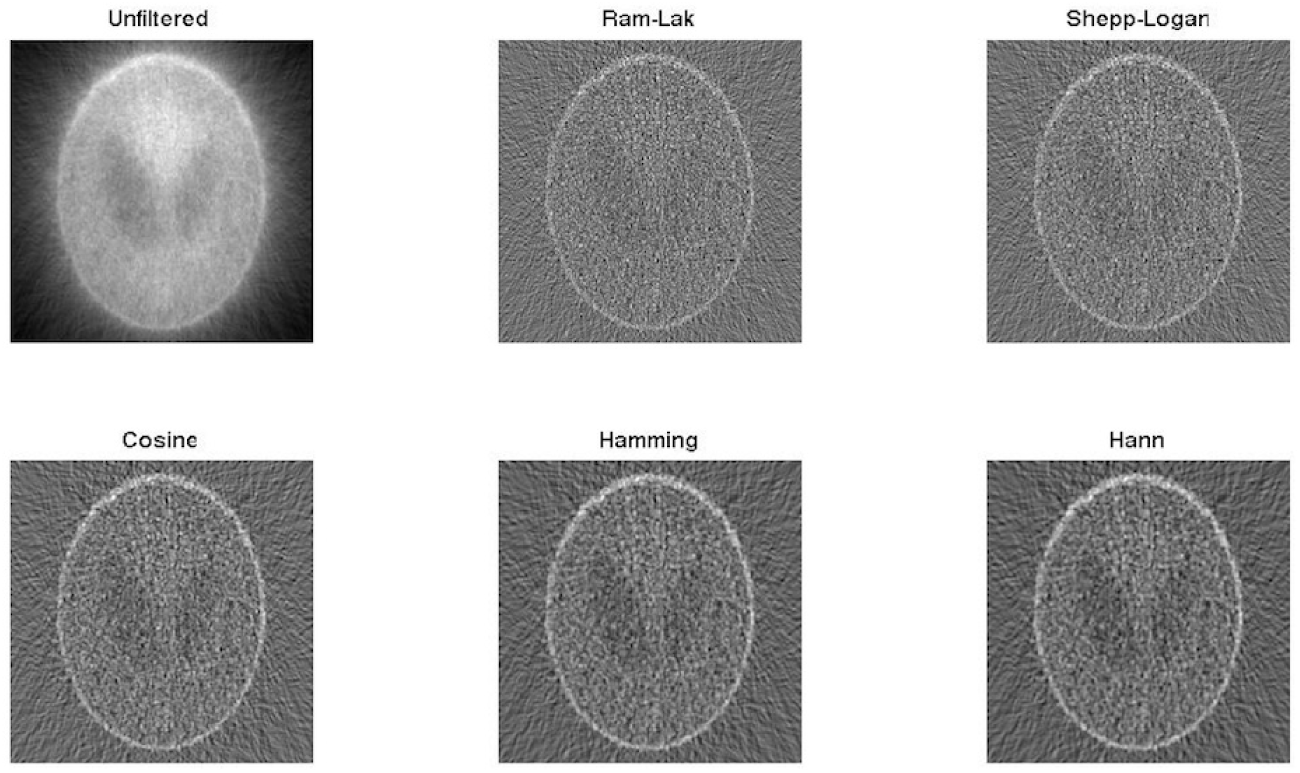
Reconstructions of the Shepp–Logan phantom, obtained from the noisy Radon sinogram (shown in the bottom part of Figure 1, panel (b)) by using the unfiltered back-projection, and the FBP algorithm with five different choices for the filtering function. Except for the case of the unfiltered back-projection, the internal structure of the phantom is almost completely lost.

While dealing with the Radon sinogram as a (rescaled) Hough sinogram, then each pixel of the noisy Radon sinogram is a cell of centre (*ϑ*_*i*_, *γ*_*j*_) in the parameter space, and the value of the pixel, multiplied by the sampling distance *d*_2_ with respect to the component *λ*_2_ = *γ* (cf. (10), for *t* = 2), is the number of straight lines characterized by parameters (*ϑ*_*i*_, *γ*_*j*_) in the image space. Note that this number need not be an integer. In fact, all pixel values in the image space 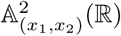 are initialized to zero and then, for any pixel centred at (*ϑ*_*i*_, *γ*_*j*_) in the parameter space 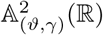 and having value *S*_*n*_(*ϑ*_*i*_, *γ*_*j*_), we trace back in 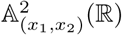 a straight line of equation *γ*_*j*_ *− x*_1_ cos *ϑ*_*i*_ *− x*_2_ sin *ϑ*_*i*_ = 0, and the value of each pixel crossed by this straight line is increased by *d*_2_*S*_*n*_(*ϑ*_*i*_, *γ*_*j*_). The resulting visualization is shown in the bottom-right panel of Figure 3. It is also interesting to implement the above procedure by taking into account only the pixels whose values are larger than a certain threshold. Various thresholds are considered, as five different percentages of the maximum value of the pixels in the noisy Radon sinogram (multiplied by *d*_2_). The corresponding visualizations are shown in the first five panels of Figure 3.

**Figure 3.**
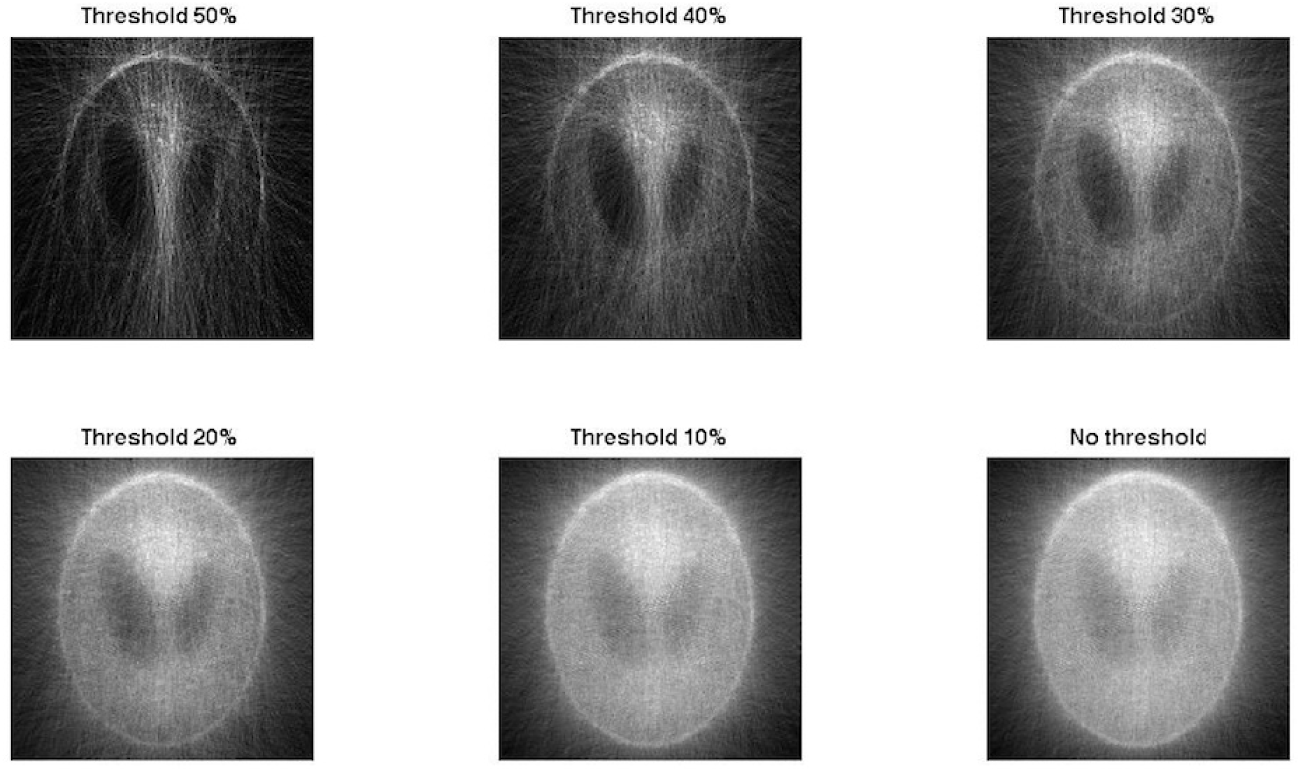
Visualizations of the Shepp–Logan phantom, obtained by drawing straight lines identified by pairs of parameters corresponding to cells in the Hough counter (i.e., the noisy Radon sinogram shown in the bottom part of Figure 1, panel (b)) whose values are higher than a fixed percentage of the maximum value. Five different thresholds are chosen, while “no threshold” means that all the pairs of parameters related to non-empty cells are used to identify straight lines in the image space.

It is worth observing that, unlike Figure 2, the pixel values in the panels of Figure 3 are not related, in principle, to the true values of the Shepp–Logan phantom.

However, a visual comparison between Figure 2 and 3 suggests that, for appropriate values of the threshold, the Hough-based approach can provide visualizations that are more informative and accurate than those provided by the (Radon-based) FBP algorithm. This is confirmed by a more quantitative analysis, where we have 1) rescaled the grey levels of all the visualizations in the range [0, 1], as in the original Shepp–Logan phantom; 2) masked the pixels of the background in order to compare just the values of the pixels inside the phantom; 3) computed the visualization error as the Frobenius norm of the matrices given by the differences between each visualization and the original Shepp–Logan phantom. The results of this analysis are in Figure 4, which shows that the Hough-based reconstruction presents a semi-convergent behavior and that it is more accurate than the Radon-based one.

**Figure 4.**
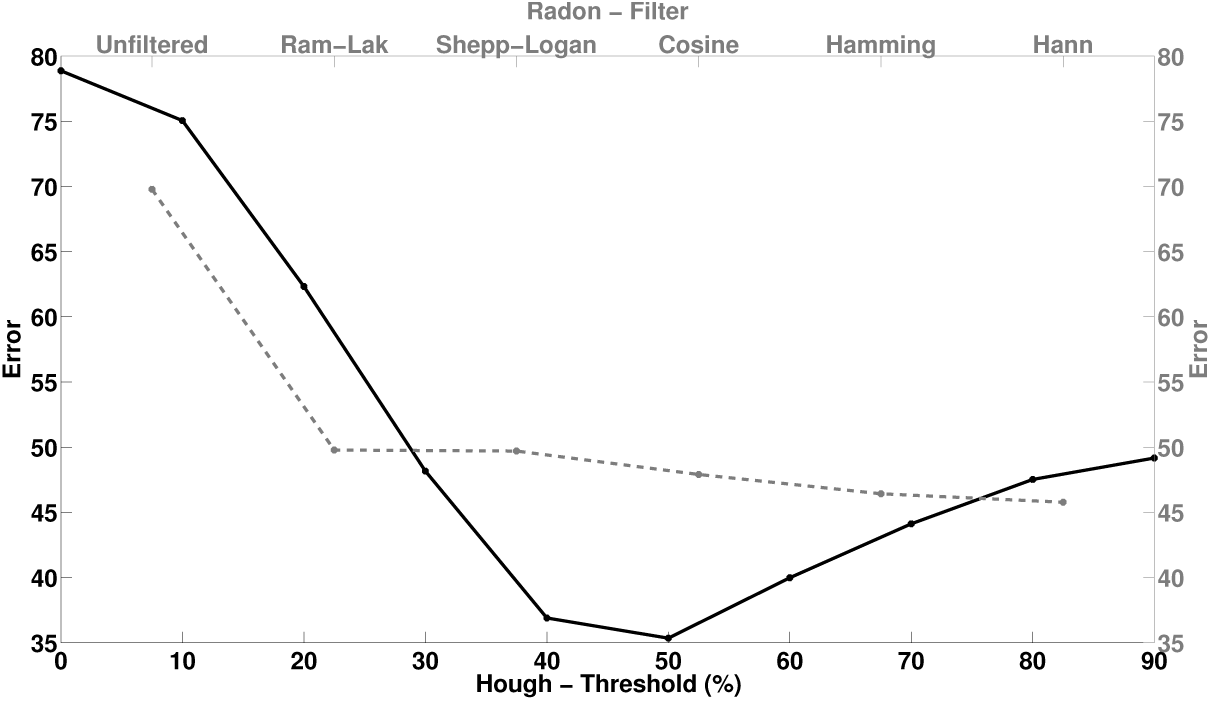
Visualization errors committed in the inversion of a noisy Radon sinogram by using unfiltered/filtered back projection with different filtering functions (grey dashed plot and grey axes) and by using the Hough-based procedure (black plot and axes).

## 5. Conclusions and future perspectives

This paper provides a rigorous description of the formal relationship between RT and HT. Specifically, the main theoretical result of the paper is concerned with the forward problem associated to RT, i.e., the proof that the rescaled Hough counter of a digital image tends to the Radon sinogram as the discretization step in the parameter space vanishes. Here a digital image is modeled by a finite linear combination of Dirac delta functions, each centered in the center of its corresponding pixel. Moreover, we briefly discussed how the relationship between RT and HT may have impacts on the inverse problem associated to image reconstruction in the case of modalities in which the data formation process is modeled by the RT. Indeed, application perspectives of this paper are concerned with the possibility to invert a Radon sinogram by regarding it as a rescaled Hough sinogram. The simple synthetic example considered in the previous section suggests that the Hough-based procedure with an optimal thershold on the Hough counter may be effectively exploited to invert noisy Radon sinograms. Further studies should be aimed at testing this computational method in the case of experimental tomographic data.

